# Outbreaks of Newcastle Disease Virus Genotype VII.1.1 in Vaccinated Commercial Poultry, Germany, 2026

**DOI:** 10.64898/2026.11.11.765530

**Authors:** Hannes Bergmann, Annett Rudovsky, Christian Grund, Ronny Marquart, Franz Kronthaler, Irene Malkwitz, Michael Winzig, Jennifer Kühn, Diana Holland, Kim S. Giessler, Lisa Rogoll, Nadine Kakerow, Jörn Gethmann, Martin Beer, Carola Sauter-Louis

**Affiliations:** Friedrich-Loeffler-Institut, Federal Research Institute for Animal Health, Institute of Epidemiology, Greifswald-Insel Riems, Germany; Friedrich-Loeffler-Institut, Federal Research Institute for Animal Health, Institute of Diagnostic Virology, Greifswald-Insel Riems, Germany; State Office for Occupational Safety, Consumer Protection and Health, Department V2, Animal Disease Prevention and Control, Potsdam, Germany; Bavarian Health and Food Safety Authority, Oberschleifiheim, Germany; Ministry of Agriculture and Food, Environment, and Consumer Protection of the State of Brandenburg, Division 6: Consumer Protection; Veterinary Services, Prevention and Control of Animal Diseases, Potsdam Germany

**Author notes:** **Correspondence:** Hannes Bergmann, Friedrich-Loeffler-Institut, Institute of Epidemiology, Südufer 10, 17493 Greifswald-Insel Riems, Germany.

**Keywords:** Newcastle disease, Newcastle disease virus, *Avian orthoavulavirus 1*, genotype VII.1.1, poultry, vaccination, surveillance, disease outbreaks, Germany

## Abstract

During February–May 2026, Newcastle disease virus subgenotype VII.1.1, related to viruses circulating in Eastern Europe, caused Germany’s first major outbreaks in decades, despite mandatory poultry vaccination programs. Clinical signs ranged from inapparent to severe, with low recorded mortality. These findings highlight the need for enhanced surveillance beyond clinical disease detection.

**Summary:** Velogenic Newcastle disease virus genotype VII.1.1 was introduced into German commercial poultry in 2026 despite mandatory vaccination; infected flocks with older birds showed low reported mortality and variable clinical signs, which challenges clinical detection-based control in the European Union.

## Introduction

Newcastle disease (ND) is caused by infection of poultry with virulent strains of *Avian orthoavulavirus 1*, also known as Newcastle disease virus (NDV), with or without clinical signs (1). Genotype VII viruses spread as a panzootic through Asia and the Middle East since about 2012 (2), causing outbreaks also in vaccinated flocks (3). Subgenotype VII.1.1 (4) spread across the Russian Federation from 2019 and was detected in Poland in 2023 (5). From 2024, it spread westward across Poland and to other European countries (6, 7).

In the European Union, ND is a category A disease, controlled by culling all animals in infected holdings and establishing restriction zones (8). The earlier infected flocks are detected, the more effectively spread and losses are limited. In Germany, vaccination against ND is mandatory for all chicken and turkey flocks (9). ND vaccination effectively prevents clinical disease, but not infection or shedding (10); VII.1.1 could therefore circulate in vaccinated flocks without typical clinical signs. On February 20, 2026, ND was confirmed in a turkey holding in Brandenburg, Germany, *<*1 km from the Polish border, after Germany has been considered free from ND for over 30 years (11). We describe the spatiotemporal pattern and the epidemiological and clinical characteristics of the ND outbreaks in Germany during February–May 2026.

## The Study

We analyzed official outbreak notifications from the EU Animal Disease Information System (ADIS) (12), based on the German animal disease notification system (Tierseuchen-Nachrichtensystem, TSN). Extended ADIS outbreak reports provided production system, vaccination history, and clinical signs for each affected holding. We excluded pigeon paramyxovirus outbreaks. Commercial holdings were classified by production system (layer, broiler, breeder, turkey) and non-commercial holdings as backyard. We identified spatial clusters by kernel density estimation. All intervals are reported relative to the date of outbreak suspicion rather than the date of laboratory confirmation. Genomes of viruses from the first outbreaks in Brandenburg and Bavaria were compared with 212 class II NDV reference genomes by neighbor-joining analysis using MEGA12 (Appendix; Appendix Table 1). Other analyses were done in R version 4.4.2.

Extended reports were available for 65 German outbreaks in domestic birds during 20 February–27 May 2026: 55 commercial flocks (28 layer, 17 broiler, 7 breeder, 3 turkey) and 10 backyard flocks. Most outbreaks (49/65, 75%) were confirmed in March (Figure 1, panel B). Genotype VII.1.1 was confirmed in all outbreaks by genotype-specific RT-qPCR. 28/65 outbreaks were classified as secondary outbreaks.

**Figure 1.**
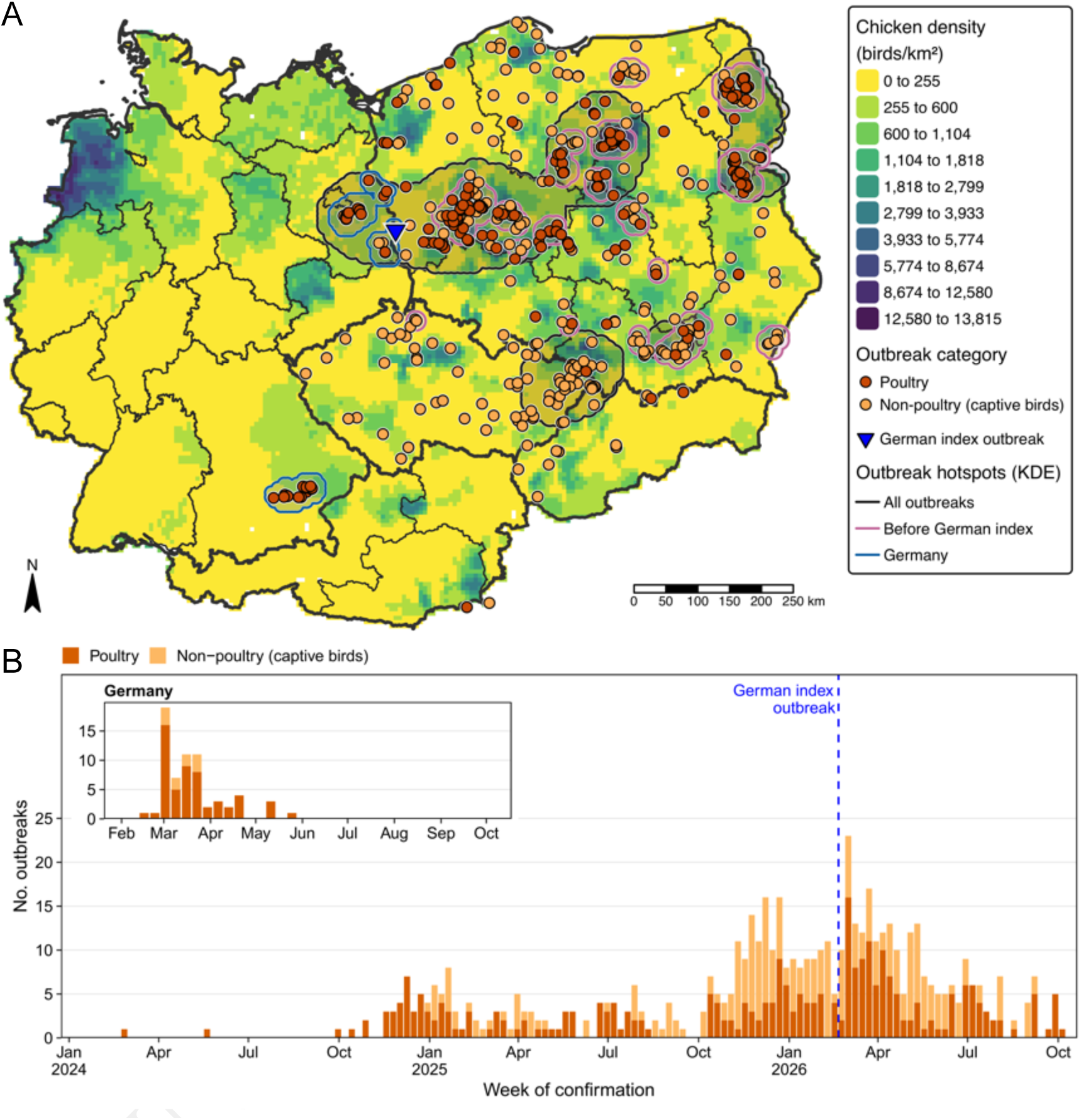
Spatial and temporal distribution of Newcastle disease (ND) outbreaks in the EU reported to the Animal Disease Information System (ADIS), January 1, 2024–October 12, 2026. A) Outbreaks in Germany and neighboring Czechia, Poland, Slovakia and Slovenia (n = 420; Germany, n = 65), by European Union disease category: vermilion, poultry; light orange, non-poultry (captive birds). The blue triangle indicates the German index outbreak, confirmed February 20, 2026. Outlined areas are kernel density estimation (KDE) hotspots, defined as areas where outbreak density exceeded the 90th (all outbreaks; dark gray; bandwidth 50 km) or 98th percentile of pixel values (outbreaks before the German index outbreak, pink, bandwidth 10 km; outbreaks in Germany, blue, bandwidth 8 km). Background shading shows chicken density (birds/km^2^, reference year 2020; Gridded Livestock of the World version 4 (GLW4), dasymetric layer, 5 arc-minute [*⇡*10 km] resolution) (13); darker shading indicates higher density. B) Weekly number of ND outbreaks in all ADIS-reporting countries (n = 551 outbreaks in 16 countries; Appendix Figure 1), by week of confirmation and disease category; the dashed line marks confirmation of the German index outbreak. Inset: outbreaks in Germany from February 2026 (n = 65). Outbreaks in wild birds and notifications in pigeons (pigeon paramyxovirus type 1) were excluded. Data sources: ADIS, accessed October 12, 2026; chicken density, Food and Agriculture Organization of the United Nations (GLW4).

Outbreaks formed 2 clusters, in Brandenburg and in Bavaria, both in areas of higher poultry density (Figure 1, panel A). The index outbreak and the Brandenburg cluster lay within the outbreak hotspot extending from Poland. Outbreak investigations identified poultry movements from infected zones in Poland to the Bavarian cluster. Genomes from both clusters (GenBank submission: SUB16616385) belonged to subgenotype VII.1.1 (4), had the polybasic F protein cleavage site motif, characteristic of virulent NDV (1), and were closely related to viruses from Eastern Europe (Figure 4).

Layers accounted for the largest proportion of outbreaks in Germany, whereas backyard holdings predominated in other European countries (Figure 2, panel A; Appendix Figure 1). All 55 commercial holdings were vaccinated against ND, as required in Germany; detailed vaccination records were un-available for 1 holding. Layers typically received 2 to 3 times an attenuated live vaccine and once an inactivated vaccine preparation during rearing, with the last dose 1 month to *>*1 year before suspicion (Figure 2, panel C). Most broilers received their last dose *≤*21 days before the suspicion, and breeder hens 1 week to 1 month before.

**Figure 2.**
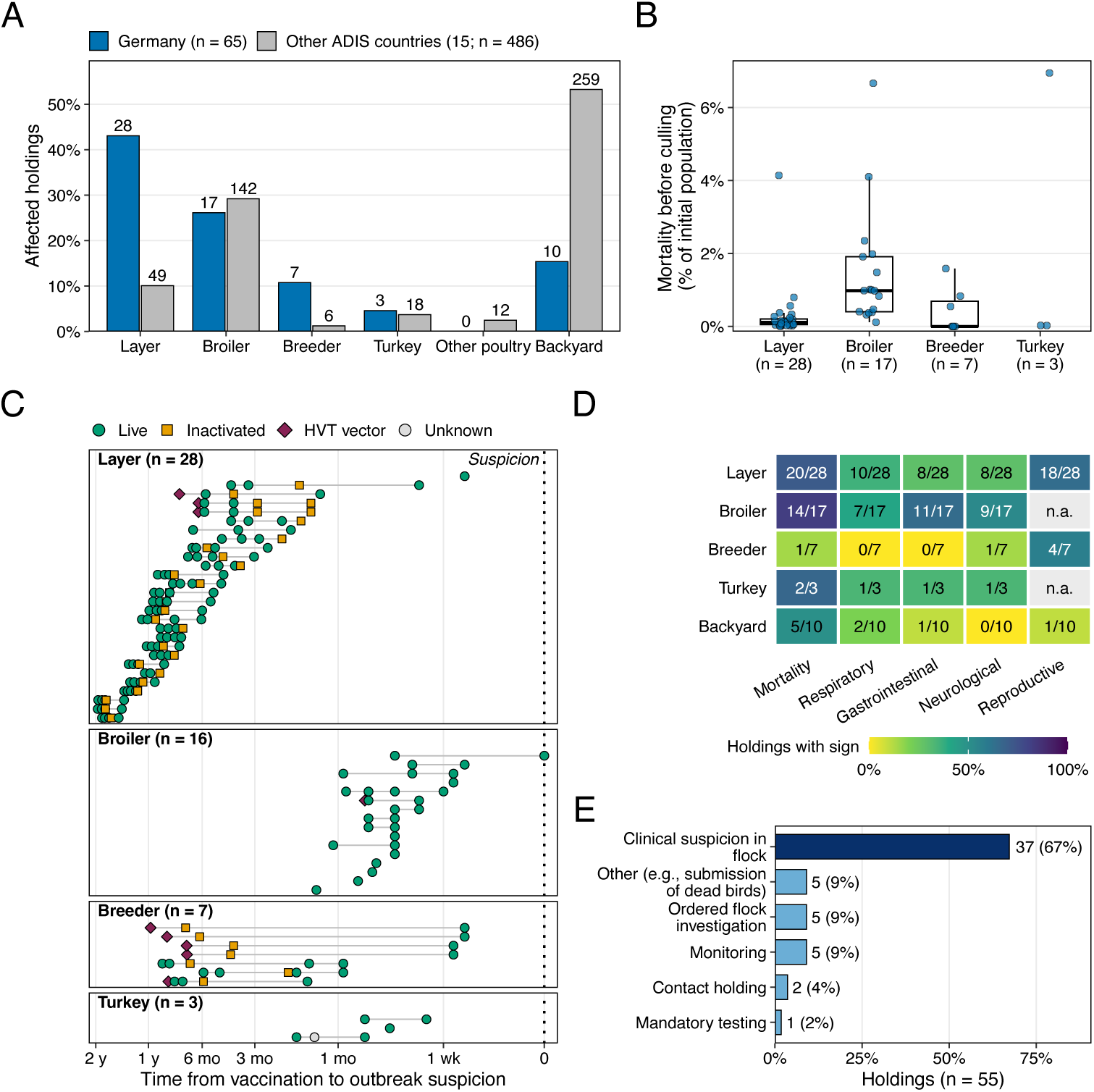
Epidemiological characteristics of Newcastle disease (ND) outbreaks in Germany and other countries reporting to the Animal Disease Information System (ADIS), 2024–2026. A) Production system of affected holdings, Germany (n = 65) and all other ADIS-reporting countries (n = 486; 15 countries; Appendix Figure 1), January 1, 2024–October 12, 2026; numbers above bars are holdings. Production system was classified from ADIS outbreak descriptions; non-poultry outbreaks are shown as backyard. B) Mortality just before culling (birds that died/initial population) in affected commercial holdings in Germany, by production system. Deaths recorded from the start of the outbreak investigation until culling; points are holdings, boxes show median and interquartile range for groups with *≥*5 holdings (Kruskal–Wallis test, *p <* 0.001); Dunn’s post hoc tests with Holm correction showed significantly higher mortality in broilers than in breeders and layers, while breeders and layers did not differ significantly. Mortality reflects only the period until culling (median 3 days after suspicion) and does not include morbidity. C) ND vaccination history of affected commercial holdings in Germany. Each row is 1 outbreak, which may include several barns. All 55 commercial holdings were vaccinated against ND, as required in Germany; detailed vaccination records were unavailable for 1 broiler holding. Points are recorded vaccine doses by type, plotted by time before outbreak suspicion (pseudo-logarithmic scale; dotted line, suspicion date). Up to 5 doses were recorded per outbreak; repeated live booster doses, such as those given to layers by drinking water or spray, are therefore not fully shown. D) Holdings with each clinical sign, by production system; values are holdings reporting the sign/all holdings of that production system, including holdings with unknown status; shading shows the proportion. n.a., not applicable (reproductive signs in meat-type birds). E) Reason for ND testing recorded in the Tierseuchen-Nachrichtensystem (TSN) for affected commercial holdings in Germany (n = 55). Panels B–E include German outbreaks with extended ADIS reports, confirmed February 20–May 27, 2026 (n = 65; commercial, n = 55). Outbreaks in wild birds and in pigeons were excluded from all panels. HVT, herpesvirus of turkeys. Data sources: ADIS (A), accessed October 12, 2026; TSN (B, E), accessed October 9, 2026, official veterinary records (C, D).

Holdings were culled at a median of 3 days (IQR 2–4) after suspicion (Appendix Figure 2). Mortality before culling differed between production systems (Kruskal–Wallis test, p*<*0.001); it was highest in broilers (median 0.98%; range 0.12–6.67%), followed by layers (median 0.11%; range 0.01–4.14%), and decreased with flock age (Figure 2, panel B; Appendix Figure 2).

Increased mortality was reported most often in layers and broilers, reproductive signs (‘egg drop’) in layers and breeders, and gastrointestinal and neurologic signs in broilers (Figure 2, panel D; Appendix Figure 3). Most holdings were detected after clinical suspicion 67% (37/55), but also through testing in restriction zones, contact tracing, or monitoring (Figure 2, panel E).

Production records showed that changes in mortality and egg production preceded suspicion, in varying order (Figure 3). On farm 1, egg production in a 60-week-old flock left the normal range on day *—*13 and daily mortality exceeded 20-fold baseline by day *—*9 (Figure 3, panels A, B). Changes started later in a 40-week-old flock on the same farm and were absent in a 101-week-old flock. On farm 3, white layers had a decline in egg production to 73% of baseline and an increase in mortality to *>*100-fold baseline within 4 days before suspicion. Brown layers in an adjacent compartment, in which infection was not confirmed, showed no change (Figure 3, panels E, F).

**Figure 3.**
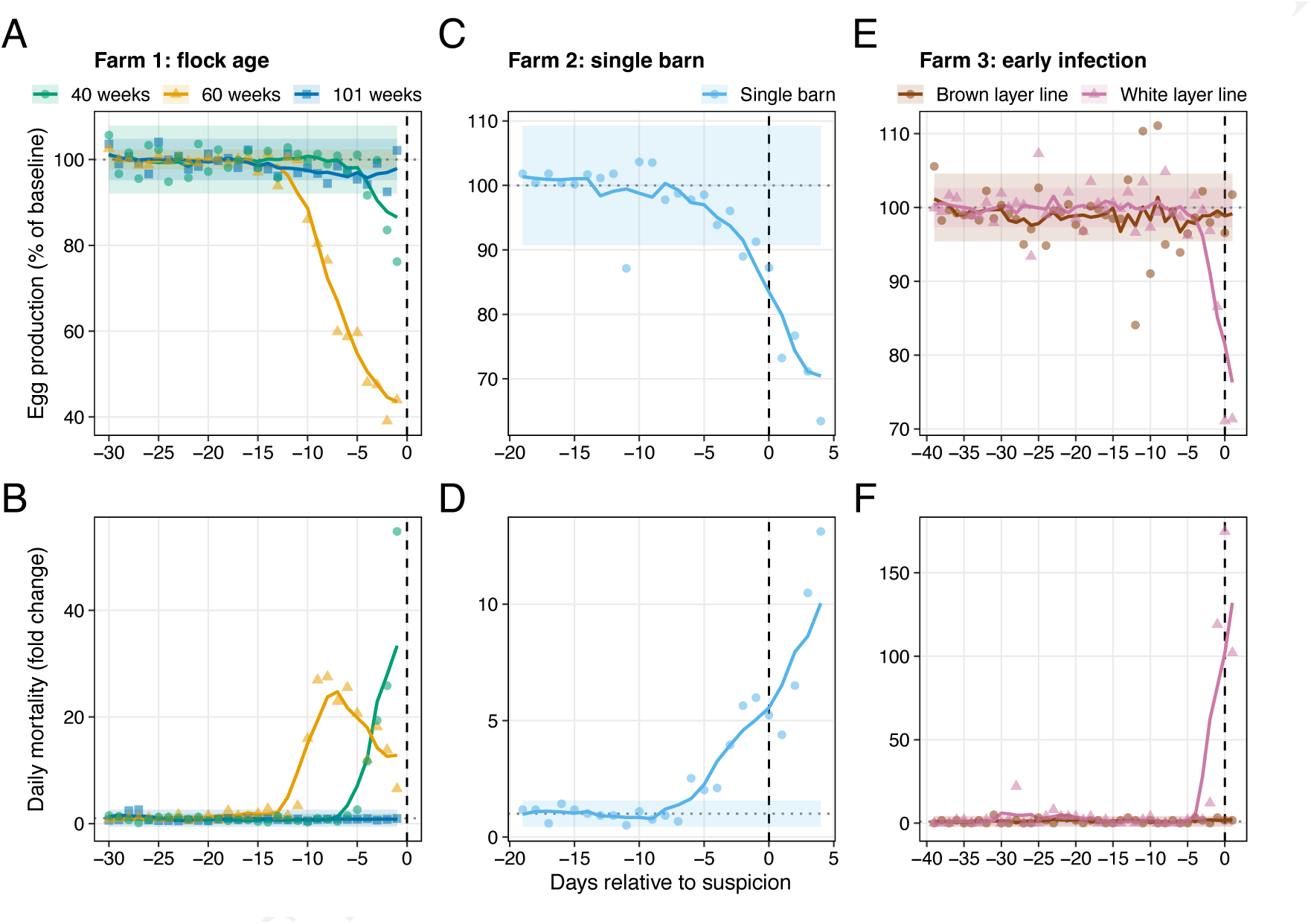
Egg production and daily mortality relative to Newcastle disease outbreak suspicion in 3 affected commercial layer farms, Germany, 2026. Columns show farms 1–3; top row, egg production (% of pre-outbreak baseline); bottom row, daily mortality (fold change from pre-outbreak baseline). Points are daily observations, lines are 5-day centered rolling means, and shaded bands show the expected normal range (baseline mean *±*2 SD; baseline, first 10 days of available data per flock). Vertical dashed line, outbreak suspicion (day 0); horizontal dotted line, baseline. A, B) Farm 1: 3 flocks aged 40, 60, and 101 weeks on the same premises. C, D) Farm 2: single barn (flock age 33 weeks). E, F) Farm 3: white and brown layers (both 49 weeks) in adjacent compartments of the same barn; infection with velogenic virus was confirmed only in the white layers. Data were collected by official veterinary authorities during outbreak investigations.

**Figure 4.**
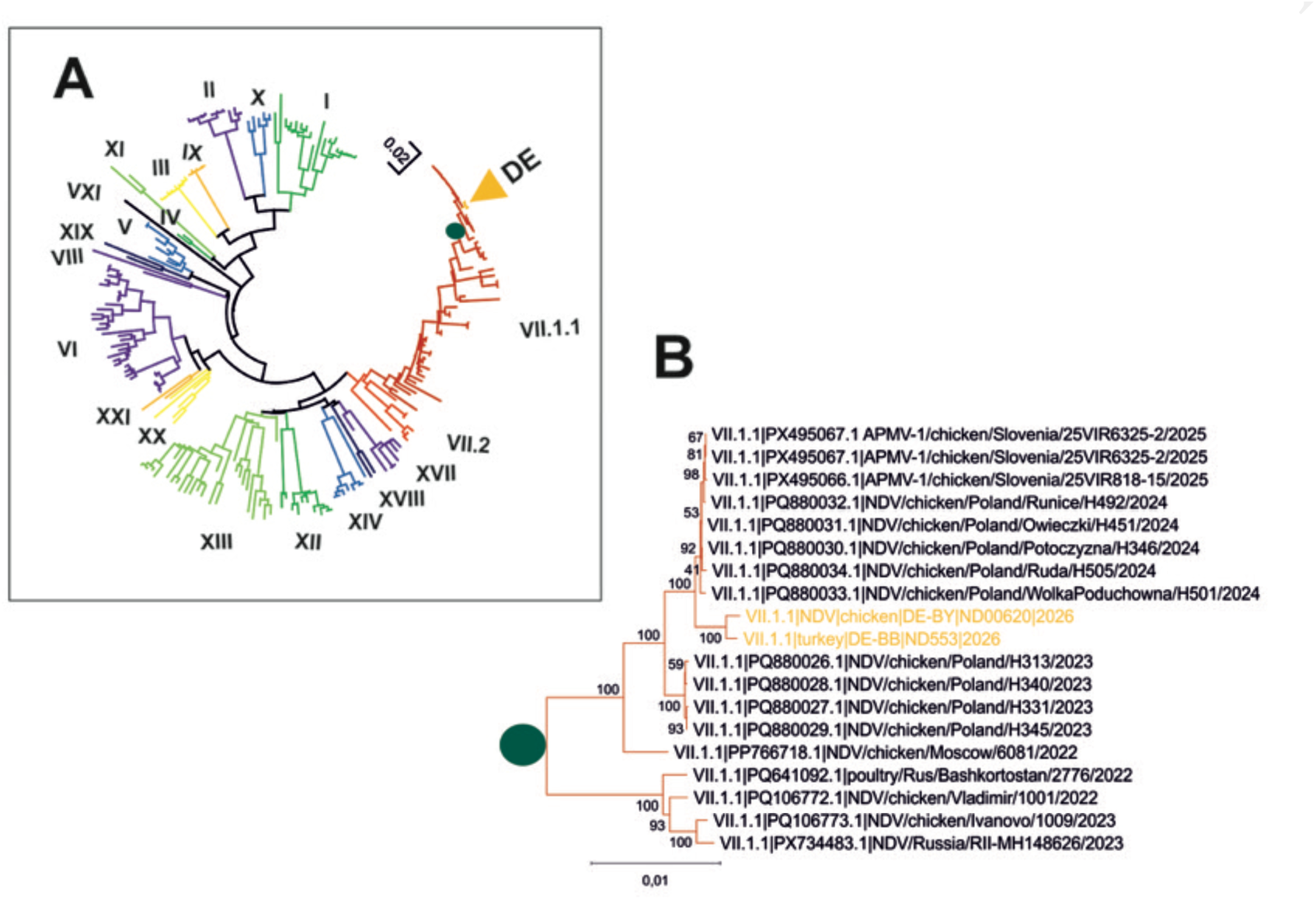
Phylogenetic analysis of selected Newcastle disease viruses from the first outbreaks in Brandenburg and Bavaria, Germany, 2026. A) Neighbor-joining tree of class II Newcastle disease viruses, including the German viruses turkey/DE-BB/ND553/2026 (Brandenburg) and chicken/DE-BY/ND620/2026 (Bavaria) and 212 reference sequences representing all recognized genotypes (4); both German viruses belong to subgenotype VII.1.1. The circle marks the root of the subtree shown in panel B and the arrowhead marks the location of the German viruses. B) Subgenotype VII.1.1 viruses from Europe; the 2 German viruses (orange labels) are closely related and cluster with viruses from Eastern Europe. Trees are based on whole-genome sequences; evolutionary distances were calculated with the Tajima–Nei model, and branch support was assessed with 1,000 bootstrap replicates. Scale bars indicate nucleotide substitutions per site. Reference sequences and GenBank accession numbers are listed in Appendix Table 1.

## Conclusions

NDV subgenotype VII.1.1 spread into commercial poultry in Germany in 2026, 30 years after the last outbreak and despite mandatory vaccination (11). Spatial clustering near the Polish border, poultry movements from Poland to the Bavarian cluster, and close genetic relatedness of German viruses to viruses from Eastern Europe are consistent with spread from the epidemic in neighboring countries (7).

Infection occurred in vaccinated holdings, often within weeks of the last booster immunization. Clinical presentation ranged from inapparent to classic neurological signs, indicating that reported mortality underestimates disease impact. Despite culling a median of 3 days after suspicion, mortality rose sharply and reached up to 6.7% in broiler flocks, while morbidity, that was frequently observed in young flocks (Appendix Figure 3), remained unquantified. These observations agree with experimental evidence that ND vaccines efficiently prevent disease, but do not induce sterile immunity, whilst still reducing infection or shedding (10, 14). Vaccine–field strain mismatch, vaccine quality or administration, and infection pressure may all have contributed but cannot be distinguished from these data (3).

In the EU and other ND-free countries, control relies on rapid detection and stamping out of infected flocks, which has eradicated previous incursions (1, 3, 8). In vaccinated populations, infected flocks may be recognized late or not at all. Monitoring mortality and egg production against flock-specific baselines, as proposed for avian influenza (15), and testing for ND after any unexplained deviation should support earlier detection. In 18 of 55 commercial holdings (33%), infection was detected not through clinical suspicion but by testing in restriction zones, tracing, or monitoring; active testing of vaccinated flocks must therefore complement clinical surveillance. As clinical signs of ND and avian influenza overlap, flocks investigated for either disease should be tested for both.

This study has limitations. Official notifications included no unvaccinated comparison group and no serologic data on flock immunity, which inform NDV transmission (2); production data came from only 3 farms, and transmission routes between holdings could not be reconstructed.

No further outbreaks were reported in Germany after May 2026, but outbreaks continued through October 2026 in neighboring countries (Figure 1), including in Poland, where ND vaccination has been mandatory since 2025 (7). Vaccination did not prevent infection or onward spread in these outbreaks, so strict biosecurity remains the primary barrier against introduction. Surveillance beyond clinical disease, evaluation of vaccine-mediated protection against infection, and reassessment of ND control strategies in vaccinated EU poultry are needed.

## Acknowledgments

We thank the veterinary authorities, farm managers, and staff for their support and cooperation during the epidemiological investigations.

## Use of artificial intelligence

Claude (Anthropic; Claude Opus 5.5) was used for language editing, style compliance and drafting R code to help with formatting the figures. All outputs were verified by the authors, who take full responsibility for the manuscript. No AI-generated images were used.

## Ethics statement

Data were collected during official investigations of natural disease outbreaks; ethics approval was not required.

## Data availability

Virus genome sequences have been submitted to GenBank (submission ID: SUB16616385) and are currently being processed. GenBank accession numbers will be provided upon release of the sequences. Other data supporting the findings of this study are available from the corresponding author upon reasonable request, subject to confidentiality requirements.

## About the Author

Dr. Bergmann is a veterinarian and immuno-epidemiology researcher at the Institute of Epidemiology, Friedrich-Loeffler-Institut, Greifswald-Insel Riems, Germany. His research focuses on understanding how the immune status of population systems shapes disease transmission dynamics and how these insights can be translated into strategic approaches to improve disease prevention and health outcomes.

## Appendix

## Virus sequencing and phylogenetic analysis

Whole genomes of 2 Newcastle disease virus isolates, from the first outbreaks in Brandenburg (turkey/DE-BB/ND553/2026) and Bavaria (chicken/DE-BY/ND620/2026), were sequenced by next-generation se-quencing as described previously (A1). The sequences covered all viral genes. Phylogenetic analyses were done in MEGA12 (A2) and included 212 reference sequences representing all recognized genotypes of class II Newcastle disease virus (Appendix Table 1), classified according to Dimitrov et al. (A3). Trees were inferred by the neighbor-joining method (A4) with evolutionary distances calculated by the Tajima–Nei model (A5); branch support was assessed by bootstrap analysis with 1,000 replicates (A6).

**Appendix Table 1.**
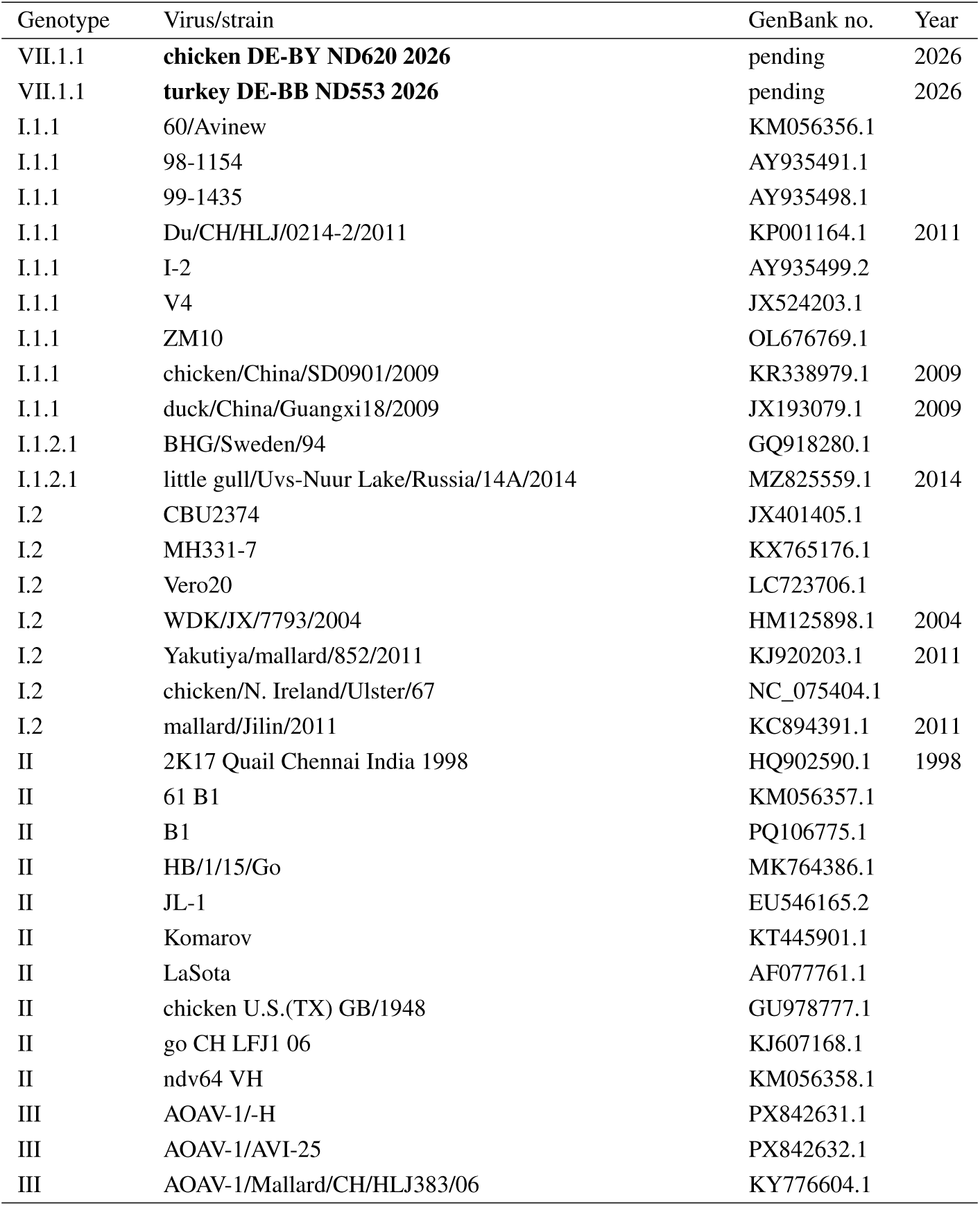
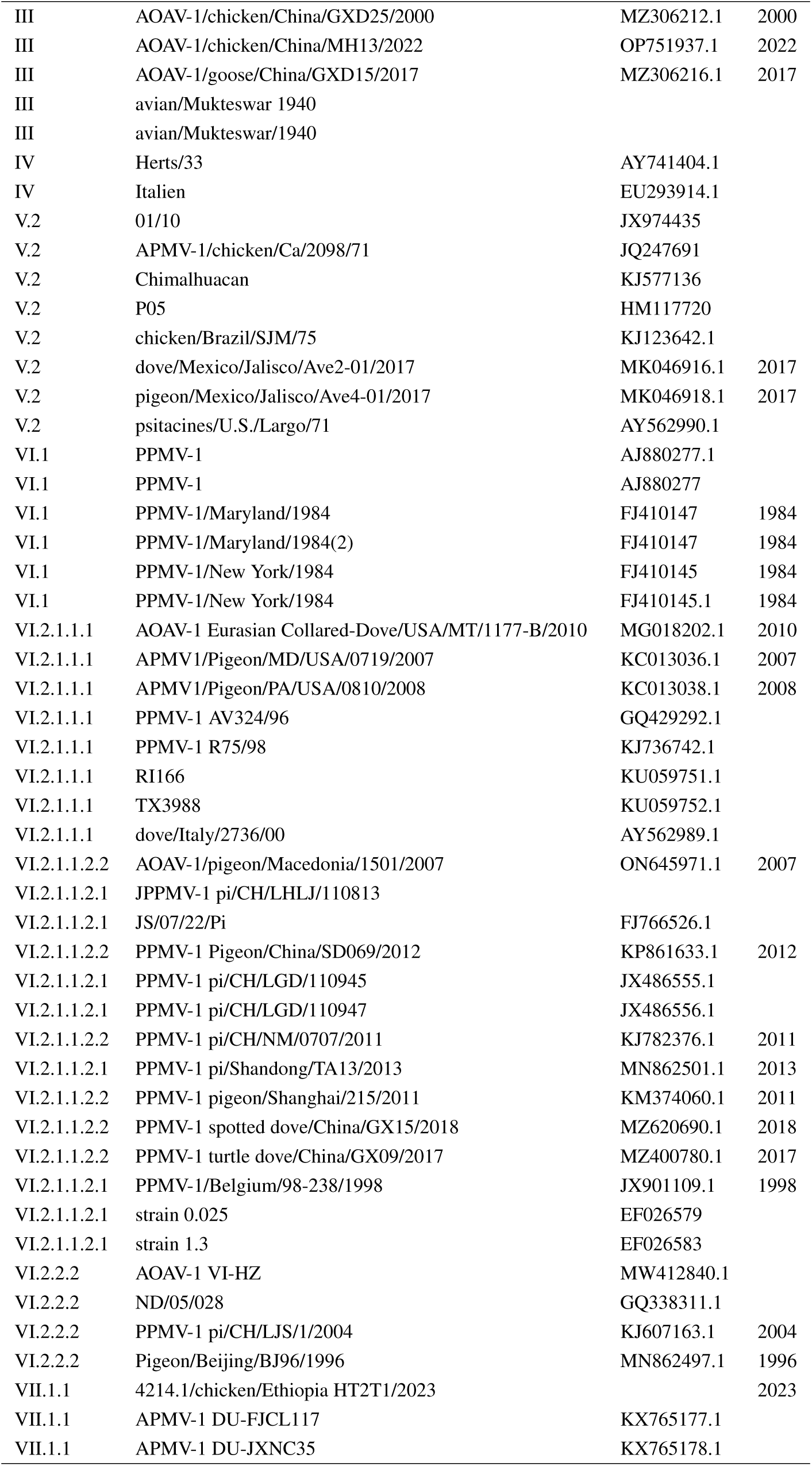
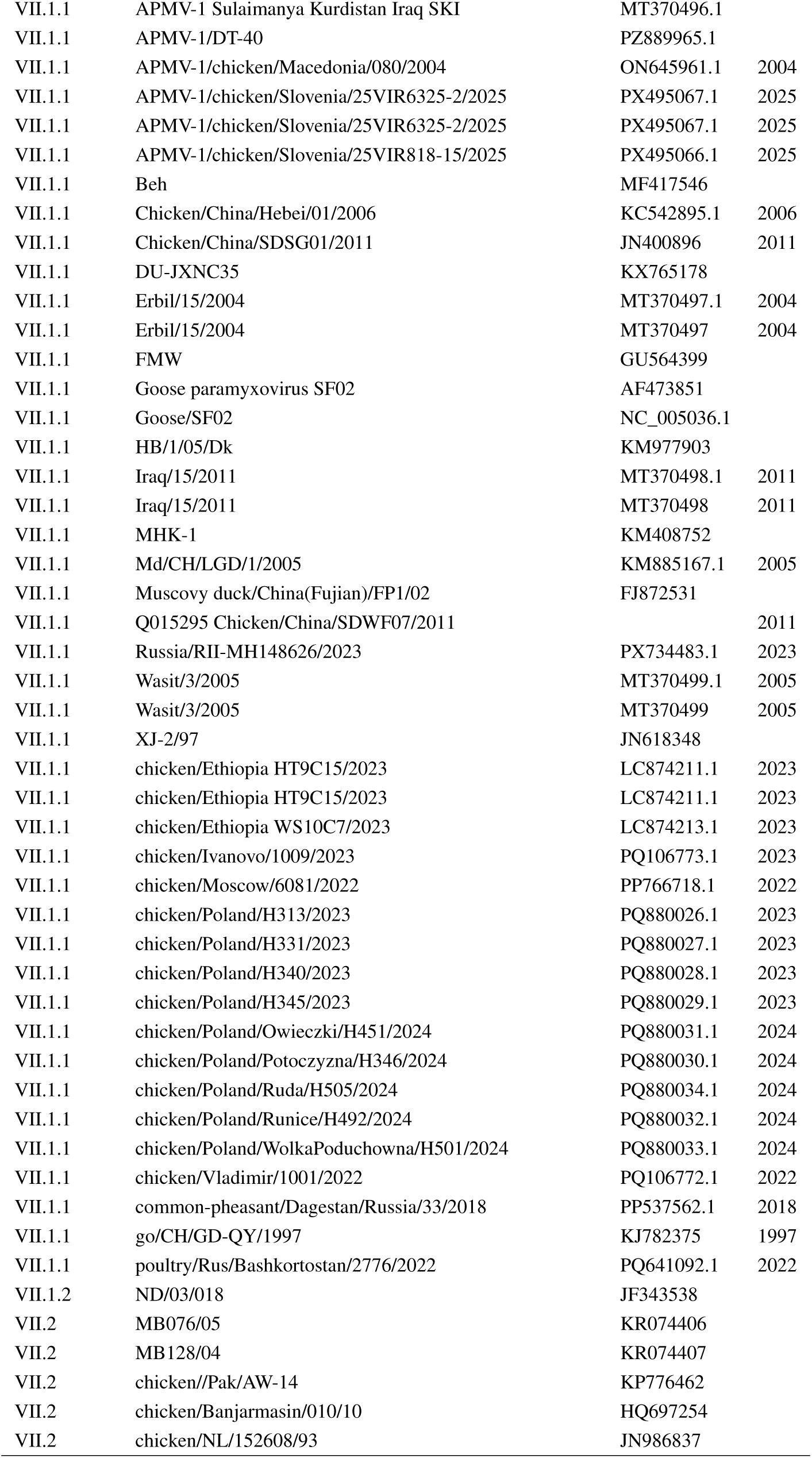
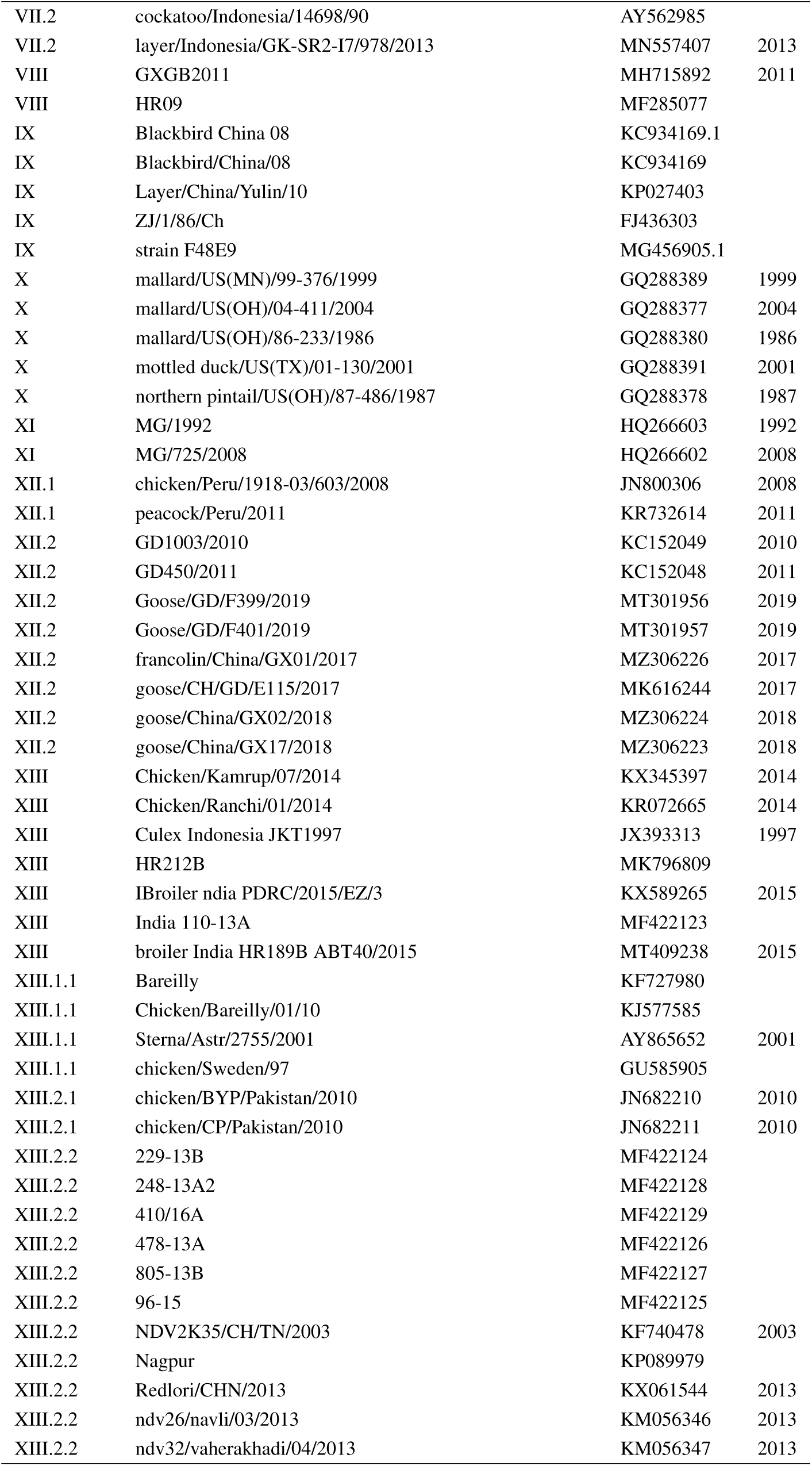
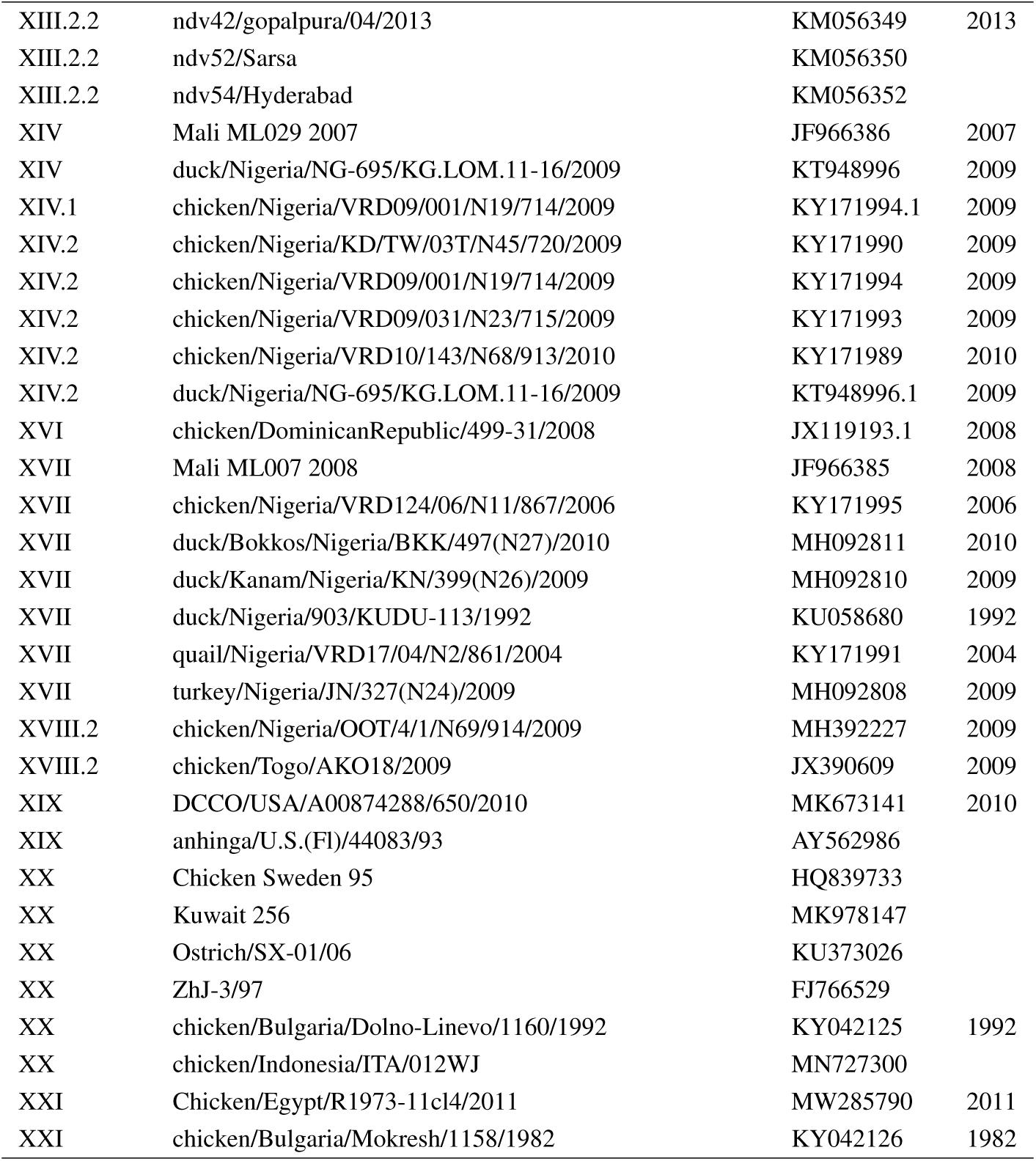
Newcastle disease virus sequences included in the phylogenetic analysis (Figure 4). Sequence labels indicate the genotypes of class 2 Newcastle disease virus (NDV) strains, followed by the host species, country of origin, isolate identification number, and year of detection. Genotype nomenclature follows the classification system proposed by Dimitrov et al. (A3), based on the fusion (F) gene sequences. For German NDV isolates (shown in bold; this study), country designations are given using ISO 3166 country codes.

| Genotype | Virus/strain | GenBank no. | Year |
|---|---|---|---|
| VII.1.1 | **chicken DE-BY ND620 2026** | pending | 2026 |
| VII.1.1 | **turkey DE-BB ND553 2026** | pending | 2026 |
| I.1.1 | 60/Avinew | KM056356.1 |  |
| I.1.1 | 98-1154 | AY935491.1 |  |
| I.1.1 | 99-1435 | AY935498.1 |  |
| I.1.1 | Du/CH/HLJ/0214-2/2011 | KP001164.1 | 2011 |
| I.1.1 | I-2 | AY935499.2 |  |
| I.1.1 | V4 | JX524203.1 |  |
| I.1.1 | ZM10 | OL676769.1 |  |
| I.1.1 | chicken/China/SD0901/2009 | KR338979.1 | 2009 |
| I.1.1 | duck/China/Guangxi18/2009 | JX193079.1 | 2009 |
| I.1.2.1 | BHG/Sweden/94 | GQ918280.1 |  |
| I.1.2.1 | little gull/Uvs-Nuur Lake/Russia/14A/2014 | MZ825559.1 | 2014 |
| I.2 | CBU2374 | JX401405.1 |  |
| I.2 | MH331-7 | KX765176.1 |  |
| I.2 | Vero20 | LC723706.1 |  |
| I.2 | WDK/JX/7793/2004 | HM125898.1 | 2004 |
| I.2 | Yakutiya/mallard/852/2011 | KJ920203.1 | 2011 |
| I.2 | chicken/N. Ireland/Ulster/67 | NC_075404.1 |  |
| I.2 | mallard/Jilin/2011 | KC894391.1 | 2011 |
| II | 2K17 Quail Chennai India 1998 | HQ902590.1 | 1998 |
| II | 61 B1 | KM056357.1 |  |
| II | B1 | PQ106775.1 |  |
| II | HB/1/15/Go | MK764386.1 |  |
| II | JL-1 | EU546165.2 |  |
| II | Komarov | KT445901.1 |  |
| II | LaSota | AF077761.1 |  |
| II | chicken U.S.(TX) GB/1948 | GU978777.1 |  |
| II | go CH LFJ1 06 | KJ607168.1 |  |
| II | ndv64 VH | KM056358.1 |  |
| III | AOAV-1/-H | PX842631.1 |  |
| III | AOAV-1/AVI-25 | PX842632.1 |  |
| III | AOAV-1/Mallard/CH/HLJ383/06 | KY776604.1 |  |
| III | AOAV-1/chicken/China/GXD25/2000 | MZ306212.1 | 2000 |
| III | AOAV-1/chicken/China/MH13/2022 | OP751937.1 | 2022 |
| III | AOAV-1/goose/China/GXD15/2017 | MZ306216.1 | 2017 |
| III | avian/Mukteswar 1940 |  |  |
| III | avian/Mukteswar/1940 |  |  |
| IV | Herts/33 | AY741404.1 |  |
| IV | Italien | EU293914.1 |  |
| V.2 | 01/10 | JX974435 |  |
| V.2 | APMV-1/chicken/Ca/2098/71 | JQ247691 |  |
| V.2 | Chimalhuacan | KJ577136 |  |
| V.2 | P05 | HM117720 |  |
| V.2 | chicken/Brazil/SJM/75 | KJ123642.1 |  |
| V.2 | dove/Mexico/Jalisco/Ave2-01/2017 | MK046916.1 | 2017 |
| V.2 | pigeon/Mexico/Jalisco/Ave4-01/2017 | MK046918.1 | 2017 |
| V.2 | psitacines/U.S./Largo/71 | AY562990.1 |  |
| VI.1 | PPMV-1 | AJ880277.1 |  |
| VI.1 | PPMV-1 | AJ880277 |  |
| VI.1 | PPMV-1/Maryland/1984 | FJ410147 | 1984 |
| VI.1 | PPMV-1/Maryland/1984(2) | FJ410147 | 1984 |
| VI.1 | PPMV-1/New York/1984 | FJ410145 | 1984 |
| VI.1 | PPMV-1/New York/1984 | FJ410145.1 | 1984 |
| VI.2.1.1.1 | AOAV-1 Eurasian Collared-Dove/USA/MT/1177-B/2010 | MG018202.1 | 2010 |
| VI.2.1.1.1 | APMV1/Pigeon/MD/USA/0719/2007 | KC013036.1 | 2007 |
| VI.2.1.1.1 | APMV1/Pigeon/PA/USA/0810/2008 | KC013038.1 | 2008 |
| VI.2.1.1.1 | PPMV-1 AV324/96 | GQ429292.1 |  |
| VI.2.1.1.1 | PPMV-1 R75/98 | KJ736742.1 |  |
| VI.2.1.1.1 | RI166 | KU059751.1 |  |
| VI.2.1.1.1 | TX3988 | KU059752.1 |  |
| VI.2.1.1.1 | dove/Italy/2736/00 | AY562989.1 |  |
| VI.2.1.1.2.2 | AOAV-1/pigeon/Macedonia/1501/2007 | ON645971.1 | 2007 |
| VI.2.1.1.2.1 | JPPMV-1 pi/CH/LHLJ/110813 |  |  |
| VI.2.1.1.2.1 | JS/07/22/Pi | FJ766526.1 |  |
| VI.2.1.1.2.2 | PPMV-1 Pigeon/China/SD069/2012 | KP861633.1 | 2012 |
| VI.2.1.1.2.1 | PPMV-1 pi/CH/LGD/110945 | JX486555.1 |  |
| VI.2.1.1.2.1 | PPMV-1 pi/CH/LGD/110947 | JX486556.1 |  |
| VI.2.1.1.2.2 | PPMV-1 pi/CH/NM/0707/2011 | KJ782376.1 | 2011 |
| VI.2.1.1.2.1 | PPMV-1 pi/Shandong/TA13/2013 | MN862501.1 | 2013 |
| VI.2.1.1.2.2 | PPMV-1 pigeon/Shanghai/215/2011 | KM374060.1 | 2011 |
| VI.2.1.1.2.2 | PPMV-1 spotted dove/China/GX15/2018 | MZ620690.1 | 2018 |
| VI.2.1.1.2.2 | PPMV-1 turtle dove/China/GX09/2017 | MZ400780.1 | 2017 |
| VI.2.1.1.2.1 | PPMV-1/Belgium/98-238/1998 | JX901109.1 | 1998 |
| VI.2.1.1.2.1 | strain 0.025 | EF026579 |  |
| VI.2.1.1.2.1 | strain 1.3 | EF026583 |  |
| VI.2.2.2 | AOAV-1 VI-HZ | MW412840.1 |  |
| VI.2.2.2 | ND/05/028 | GQ338311.1 |  |
| VI.2.2.2 | PPMV-1 pi/CH/LJS/1/2004 | KJ607163.1 | 2004 |
| VI.2.2.2 | Pigeon/Beijing/BJ96/1996 | MN862497.1 | 1996 |
| VII.1.1 | 4214.1/chicken/Ethiopia HT2T1/2023 |  | 2023 |
| VII.1.1 | APMV-1 DU-FJCL117 | KX765177.1 |  |
| VII.1.1 | APMV-1 DU-JXNC35 | KX765178.1 |  |
| VII.1.1 | APMV-1 Sulaimanya Kurdistan Iraq SKI | MT370496.1 |  |
| VII.1.1 | APMV-1/DT-40 | PZ889965.1 |  |
| VII.1.1 | APMV-1/chicken/Macedonia/080/2004 | ON645961.1 | 2004 |
| VII.1.1 | APMV-1/chicken/Slovenia/25VIR6325-2/2025 | PX495067.1 | 2025 |
| VII.1.1 | APMV-1/chicken/Slovenia/25VIR6325-2/2025 | PX495067.1 | 2025 |
| VII.1.1 | APMV-1/chicken/Slovenia/25VIR818-15/2025 | PX495066.1 | 2025 |
| VII.1.1 | Beh | MF417546 |  |
| VII.1.1 | Chicken/China/Hebei/01/2006 | KC542895.1 | 2006 |
| VII.1.1 | Chicken/China/SDSG01/2011 | JN400896 | 2011 |
| VII.1.1 | DU-JXNC35 | KX765178 |  |
| VII.1.1 | Erbil/15/2004 | MT370497.1 | 2004 |
| VII.1.1 | Erbil/15/2004 | MT370497 | 2004 |
| VII.1.1 | FMW | GU564399 |  |
| VII.1.1 | Goose paramyxovirus SF02 | AF473851 |  |
| VII.1.1 | Goose/SF02 | NC_005036.1 |  |
| VII.1.1 | HB/1/05/Dk | KM977903 |  |
| VII.1.1 | Iraq/15/2011 | MT370498.1 | 2011 |
| VII.1.1 | Iraq/15/2011 | MT370498 | 2011 |
| VII.1.1 | MHK-1 | KM408752 |  |
| VII.1.1 | Md/CH/LGD/1/2005 | KM885167.1 | 2005 |
| VII.1.1 | Muscovy duck/China(Fujian)/FP1/02 | FJ872531 |  |
| VII.1.1 | Q015295 Chicken/China/SDWF07/2011 |  | 2011 |
| VII.1.1 | Russia/RII-MH148626/2023 | PX734483.1 | 2023 |
| VII.1.1 | Wasit/3/2005 | MT370499.1 | 2005 |
| VII.1.1 | Wasit/3/2005 | MT370499 | 2005 |
| VII.1.1 | XJ-2/97 | JN618348 |  |
| VII.1.1 | chicken/Ethiopia HT9C15/2023 | LC874211.1 | 2023 |
| VII.1.1 | chicken/Ethiopia HT9C15/2023 | LC874211.1 | 2023 |
| VII.1.1 | chicken/Ethiopia WS10C7/2023 | LC874213.1 | 2023 |
| VII.1.1 | chicken/Ivanovo/1009/2023 | PQ106773.1 | 2023 |
| VII.1.1 | chicken/Moscow/6081/2022 | PP766718.1 | 2022 |
| VII.1.1 | chicken/Poland/H313/2023 | PQ880026.1 | 2023 |
| VII.1.1 | chicken/Poland/H331/2023 | PQ880027.1 | 2023 |
| VII.1.1 | chicken/Poland/H340/2023 | PQ880028.1 | 2023 |
| VII.1.1 | chicken/Poland/H345/2023 | PQ880029.1 | 2023 |
| VII.1.1 | chicken/Poland/Owieczki/H451/2024 | PQ880031.1 | 2024 |
| VII.1.1 | chicken/Poland/Potoczyzna/H346/2024 | PQ880030.1 | 2024 |
| VII.1.1 | chicken/Poland/Ruda/H505/2024 | PQ880034.1 | 2024 |
| VII.1.1 | chicken/Poland/Runice/H492/2024 | PQ880032.1 | 2024 |
| VII.1.1 | chicken/Poland/WolkaPoduchowna/H501/2024 | PQ880033.1 | 2024 |
| VII.1.1 | chicken/Vladimir/1001/2022 | PQ106772.1 | 2022 |
| VII.1.1 | common-pheasant/Dagestan/Russia/33/2018 | PP537562.1 | 2018 |
| VII.1.1 | go/CH/GD-QY/1997 | KJ782375 | 1997 |
| VII.1.1 | poultry/Rus/Bashkortostan/2776/2022 | PQ641092.1 | 2022 |
| VII.1.2 | ND/03/018 | JF343538 |  |
| VII.2 | MB076/05 | KR074406 |  |
| VII.2 | MB128/04 | KR074407 |  |
| VII.2 | chicken//Pak/AW-14 | KP776462 |  |
| VII.2 | chicken/Banjarmasin/010/10 | HQ697254 |  |
| VII.2 | chicken/NL/152608/93 | JN986837 |  |
| VII.2 | cockatoo/Indonesia/14698/90 | AY562985 |  |
| VII.2 | layer/Indonesia/GK-SR2-I7/978/2013 | MN557407 | 2013 |
| VIII | GXGB2011 | MH715892 | 2011 |
| VIII | HR09 | MF285077 |  |
| IX | Blackbird China 08 | KC934169.1 |  |
| IX | Blackbird/China/08 | KC934169 |  |
| IX | Layer/China/Yulin/10 | KP027403 |  |
| IX | ZJ/1/86/Ch | FJ436303 |  |
| IX | strain F48E9 | MG456905.1 |  |
| X | mallard/US(MN)/99-376/1999 | GQ288389 | 1999 |
| X | mallard/US(OH)/04-411/2004 | GQ288377 | 2004 |
| X | mallard/US(OH)/86-233/1986 | GQ288380 | 1986 |
| X | mottled duck/US(TX)/01-130/2001 | GQ288391 | 2001 |
| X | northern pintail/US(OH)/87-486/1987 | GQ288378 | 1987 |
| XI | MG/1992 | HQ266603 | 1992 |
| XI | MG/725/2008 | HQ266602 | 2008 |
| XII.1 | chicken/Peru/1918-03/603/2008 | JN800306 | 2008 |
| XII.1 | peacock/Peru/2011 | KR732614 | 2011 |
| XII.2 | GD1003/2010 | KC152049 | 2010 |
| XII.2 | GD450/2011 | KC152048 | 2011 |
| XII.2 | Goose/GD/F399/2019 | MT301956 | 2019 |
| XII.2 | Goose/GD/F401/2019 | MT301957 | 2019 |
| XII.2 | francolin/China/GX01/2017 | MZ306226 | 2017 |
| XII.2 | goose/CH/GD/E115/2017 | MK616244 | 2017 |
| XII.2 | goose/China/GX02/2018 | MZ306224 | 2018 |
| XII.2 | goose/China/GX17/2018 | MZ306223 | 2018 |
| XIII | Chicken/Kamrup/07/2014 | KX345397 | 2014 |
| XIII | Chicken/Ranchi/01/2014 | KR072665 | 2014 |
| XIII | Culex Indonesia JKT1997 | JX393313 | 1997 |
| XIII | HR212B | MK796809 |  |
| XIII | IBroiler ndia PDRC/2015/EZ/3 | KX589265 | 2015 |
| XIII | India 110-13A | MF422123 |  |
| XIII | broiler India HR189B ABT40/2015 | MT409238 | 2015 |
| XIII.1.1 | Bareilly | KF727980 |  |
| XIII.1.1 | Chicken/Bareilly/01/10 | KJ577585 |  |
| XIII.1.1 | Sterna/Astr/2755/2001 | AY865652 | 2001 |
| XIII.1.1 | chicken/Sweden/97 | GU585905 |  |
| XIII.2.1 | chicken/BYP/Pakistan/2010 | JN682210 | 2010 |
| XIII.2.1 | chicken/CP/Pakistan/2010 | JN682211 | 2010 |
| XIII.2.2 | 229-13B | MF422124 |  |
| XIII.2.2 | 248-13A2 | MF422128 |  |
| XIII.2.2 | 410/16A | MF422129 |  |
| XIII.2.2 | 478-13A | MF422126 |  |
| XIII.2.2 | 805-13B | MF422127 |  |
| XIII.2.2 | 96-15 | MF422125 |  |
| XIII.2.2 | NDV2K35/CH/TN/2003 | KF740478 | 2003 |
| XIII.2.2 | Nagpur | KP089979 |  |
| XIII.2.2 | Redlori/CHN/2013 | KX061544 | 2013 |
| XIII.2.2 | ndv26/navli/03/2013 | KM056346 | 2013 |
| XIII.2.2 | ndv32/vaherakhadi/04/2013 | KM056347 | 2013 |
| XIII.2.2 | ndv42/gopalpura/04/2013 | KM056349 | 2013 |
| XIII.2.2 | ndv52/Sarsa | KM056350 |  |
| XIII.2.2 | ndv54/Hyderabad | KM056352 |  |
| XIV | Mali ML029 2007 | JF966386 | 2007 |
| XIV | duck/Nigeria/NG-695/KG.LOM.11-16/2009 | KT948996 | 2009 |
| XIV.1 | chicken/Nigeria/VRD09/001/N19/714/2009 | KY171994.1 | 2009 |
| XIV.2 | chicken/Nigeria/KD/TW/03T/N45/720/2009 | KY171990 | 2009 |
| XIV.2 | chicken/Nigeria/VRD09/001/N19/714/2009 | KY171994 | 2009 |
| XIV.2 | chicken/Nigeria/VRD09/031/N23/715/2009 | KY171993 | 2009 |
| XIV.2 | chicken/Nigeria/VRD10/143/N68/913/2010 | KY171989 | 2010 |
| XIV.2 | duck/Nigeria/NG-695/KG.LOM.11-16/2009 | KT948996.1 | 2009 |
| XVI | chicken/DominicanRepublic/499-31/2008 | JX119193.1 | 2008 |
| XVII | Mali ML007 2008 | JF966385 | 2008 |
| XVII | chicken/Nigeria/VRD124/06/N11/867/2006 | KY171995 | 2006 |
| XVII | duck/Bokkos/Nigeria/BKK/497(N27)/2010 | MH092811 | 2010 |
| XVII | duck/Kanam/Nigeria/KN/399(N26)/2009 | MH092810 | 2009 |
| XVII | duck/Nigeria/903/KUDU-113/1992 | KU058680 | 1992 |
| XVII | quail/Nigeria/VRD17/04/N2/861/2004 | KY171991 | 2004 |
| XVII | turkey/Nigeria/JN/327(N24)/2009 | MH092808 | 2009 |
| XVIII.2 | chicken/Nigeria/OOT/4/1/N69/914/2009 | MH392227 | 2009 |
| XVIII.2 | chicken/Togo/AKO18/2009 | JX390609 | 2009 |
| XIX | DCCO/USA/A00874288/650/2010 | MK673141 | 2010 |
| XIX | anhinga/U.S.(Fl)/44083/93 | AY562986 |  |
| XX | Chicken Sweden 95 | HQ839733 |  |
| XX | Kuwait 256 | MK978147 |  |
| XX | Ostrich/SX-01/06 | KU373026 |  |
| XX | ZhJ-3/97 | FJ766529 |  |
| XX | chicken/Bulgaria/Dolno-Linevo/1160/1992 | KY042125 | 1992 |
| XX | chicken/Indonesia/ITA/012WJ | MN727300 |  |
| XXI | Chicken/Egypt/R1973-11cl4/2011 | MW285790 | 2011 |
| XXI | chicken/Bulgaria/Mokresh/1158/1982 | KY042126 | 1982 |

---

**Appendix Figure 1.**
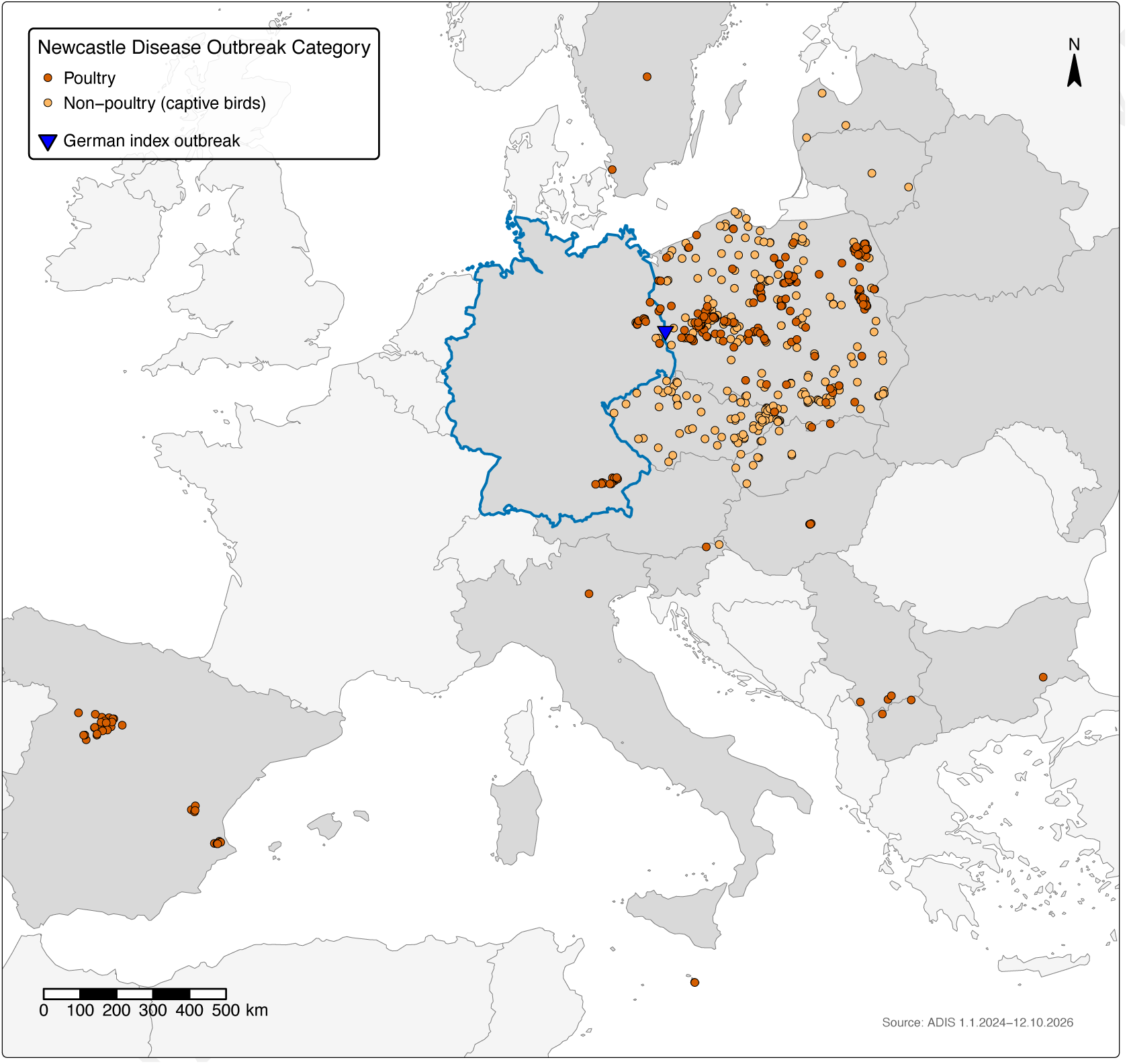
Newcastle disease outbreaks in the EU reported to the Animal Disease Information System (ADIS) by all reporting countries, January 1, 2024–October 12, 2026. The map shows the outbreaks included in Figure 1, panel B, and Figure 2, panel A (n = 551 total; 16 countries), by European Union disease category: vermilion, poultry; light orange, non-poultry (captive birds). The blue triangle indicates the German index outbreak; Germany is outlined in blue. Countries reporting *≥*1 outbreak are shaded darker. Outbreaks in wild birds and notifications in pigeons were excluded. 1 outbreak outside the map extent is not shown. Projection: ETRS89 Lambert azimuthal equal-area (EPSG:3035). Administrative boundaries: © EuroGeographics (GISCO, Eurostat).

**Appendix Figure 2.**
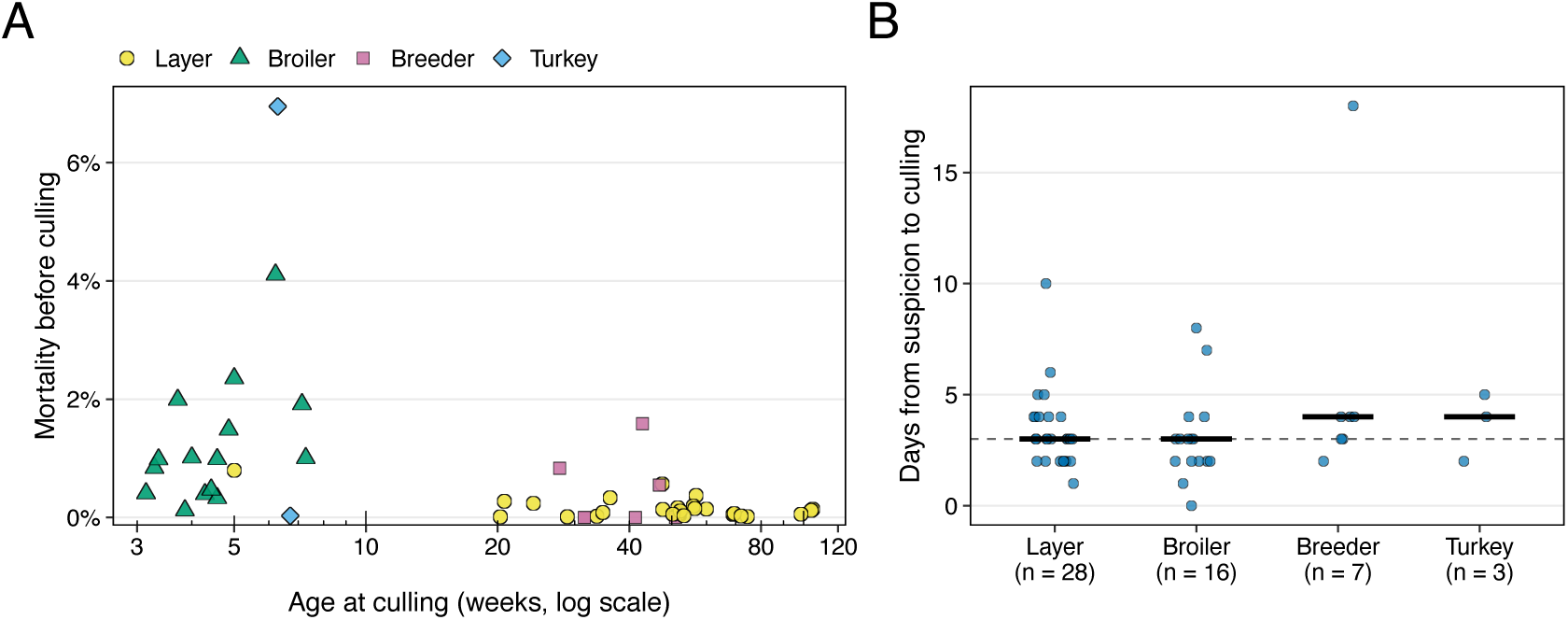
Mortality by flock age and time to culling in commercial poultry holdings affected by Newcastle disease, Germany, 2026. A) Mortality before culling (birds that died/initial population) by age at culling (weeks, logarithmic scale) and production system (n = 51 holdings with available age records; Spearman *r* = *—*0.53, *p <* 0.001). Age at culling was calculated from the age reported in extended ADIS outbreak reports and the culling date recorded in TSN. B) Days from outbreak suspicion to culling, by production system; points are holdings, horizontal bars show medians, and the dashed line shows the overall median (3 days; n = 54 holdings with available cull dates). Suspicion date was used because confirmation was sometimes recorded after culling had started. ADIS, Animal Disease Information System; TSN, Tierseuchen-Nachrichtensystem. Data sources: ADIS, accessed October 12, 2026; TSN, accessed October 9, 2026.

**Appendix Figure 3.**
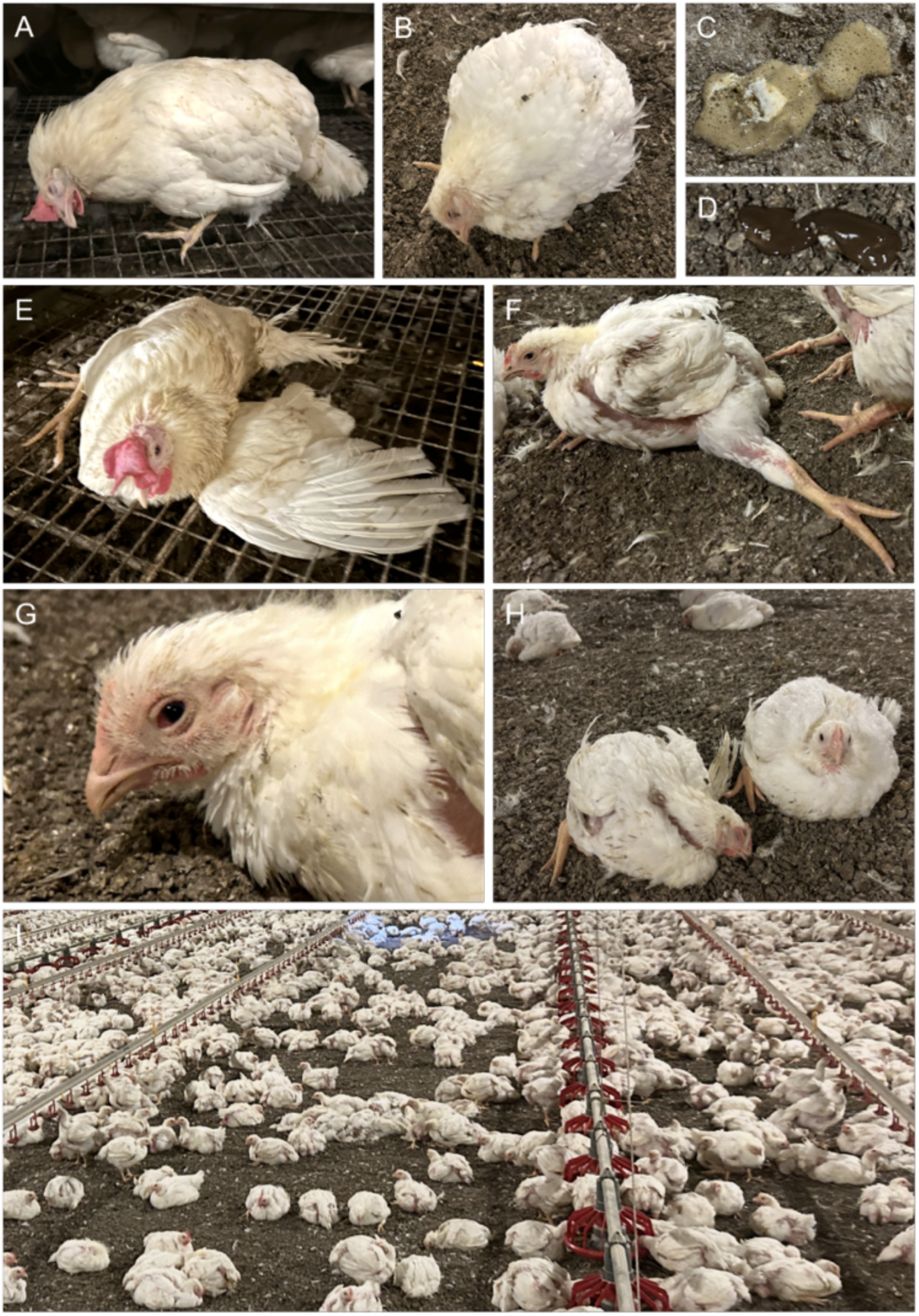
Clinical signs of Newcastle disease in affected commercial poultry flocks, Germany, 2026. A) Adult white layer hen with torticollis; the head is bent ventrally. B) Broiler chicken with torticollis, ruffled feathers, and pale skin. C) Foamy, watery, light-brown to greenish diarrhea in a broiler breeder flock without other overt clinical signs. D) Dark brown viscous diarrhea without urates in an affected broiler flock. E) White layer bird recumbent on the slatted floor of a cage-free system, with legs displaced laterally, wing and tail extended, and mild head torsion. F) Broiler chicken with splayed legs. G) Head of a broiler chicken showing hyperaemia and congestion of the nictitating membrane. H) Broiler chickens in respiratory distress, sitting with ruffled feathers, beaks resting on the litter, and eyes closed. I) Affected broiler house showing uneven clustering of birds, ruffled plumage, and reduced activity. All flocks shown were vaccinated against Newcastle disease. Images were taken by the authors during official outbreak investigations.

